# Threat of mining to African great apes

**DOI:** 10.1101/2023.10.17.562472

**Authors:** Jessica Junker, Luise Quoss, Jose Valdez, Mimi Arandjelovic, Abdulai Barrie, Genevieve Campbell, Stefanie Heinicke, Tatyana Humle, Célestin Yao Kouakou, Hjalmar S. Kühl, Isabel Ordaz-Nemeth, Henrique M. Pereira, Helga Rainer, Johannes Refisch, Laura Sonter, Tenekwetche Sop

**Affiliations:** Institute of Biology, Martin Luther University Halle-Wittenberg, Am Kirchtor 1, 06108 Halle, Germany; German Centre for Integrative Biodiversity Research (iDiv) Halle-Jena-Leipzig, Puschstrasse 4, 04103 Leipzig, Germany; Max-Planck Institute for Evolutionary Anthropology, Department of Primate Behavior and Evolution, Deutscher Platz 6, 04103 Leipzig, Germany; Ministry of Environment and Climate Change, 55 Wilkinson Road, Freetown, Sierra Leone; Re: wild, 500 N Capital of Texas Hwy Building 1, Suite 200, Austin, TX 78746, United States; Potsdam Institute for Climate Impact Research (PIK), Member of the Leibniz Association, Potsdam, Germany; Durrell of Institute of Conservation and Ecology, School of Anthropology and Conservation, University of Kent, Canterbury CT2 7NR, United Kingdom; Université Jean Lorougnon Guédé, BP 150 Daloa, Côte d’Ivoire; Senckenberg Museum for Natural History Görlitz, Senckenberg – Member of the Leibniz Association Am Museum 1, 02826 Görlitz, Germany; International Institute Zittau, Technische Universität Dresden, Markt 23, 02763 Zittau, Germany; CIBIO, Centro de Investigação em Biodiversidade e Recursos Genéticos, InBIO Laboratório Associado, Campus de Vairão, Universidade do Porto, 4485-661 Vairão, Portugal; Independent consultant, PO Box 4107, Kampala, Uganda; Great Apes Survival Partnership, United Nations Environment Programme, P.O. Box 30552 (00100), Nairobi, Kenya; School of the Environment, The University of Queensland, St Lucia 4072, Australia; Centre for Biodiversity and Conservation Science, The University of Queensland, St Lucia 4072, Australia; Sustainable Minerals Institute, The University of Queensland, St Lucia 4072, Australia

**Author notes:** Corresponding author’s. These authors contributed equally to this work (Jessica Junker, Luise Quoss, Jose Valdez).

## Abstract

The rapid growth of clean energy technologies is driving a rising demand for critical minerals. In 2022 at the UN Biodiversity Conference (COP 15), seven major economies formed an alliance to enhance the sustainability of mining these essential decarbonization minerals. However, there is a scarcity of studies assessing the threat of mining to global biodiversity. By integrating a global mining dataset with ape density distribution estimates, we explored the potential negative impact of industrial mining on African great apes. Our findings reveal that up to one-third of Africa’s great ape population faces mining-related risks. This is especially pronounced in West Africa, where numerous mining areas overlap with fragmented ape habitats, often occurring in high-density ape regions. For 97% of mining areas, no ape survey data are available, underscoring the importance of increased accessibility to environmental data within the mining sector to facilitate research into the complex interactions between mining, climate, biodiversity and sustainability.

**Teaser:** Mining for clean energy minerals could put one-third of Africa’s ape population at risk.

## Introduction

Africa is experiencing an unprecedented mining boom (*1*) threatening wildlife populations and whole ecosystems. Mining activities are growing in intensity and scale, and with increasing exploration and production in previously unexploited areas. Africa contains around 30% of the world’s mineral resources, yet less than 5% of the global mineral exploitation has occurred in Africa, highlighting the enormous potential for growth in this sector (*1*). Significant production increases in the renewable energy sector are expected to cause a boom in mineral exploitation (*2*). Africa, which is rich in ecological diversity, harbors around one-sixth of the world’s remaining forests and is home to one quarter of the world’s mammal species (*3*). Among these are primates, which are one of the most threatened groups of species, with 67% of all primate species (Africa: 73.1%) currently listed as threatened by the International Union for Conservation of Nature’s (IUCN) Red List and 42% with continuing declining population trends (*4*). Great apes (hereafter ‘apes’) are particularly at risk, with all 14 taxa currently listed as either Endangered (EN) or Critically Endangered (CR) (*5*).

Apes are our closest evolutionary relatives and are important in many societies, integral to economic and cultural wellbeing. They generate substantial income from tourism projects, and serve as powerful flagship species due to their anthropological significance, helping to raise public awareness and millions in conservation spending (*6*). They fulfill the important role of umbrella species implying that if conservation efforts focus on ape populations and their habitats, this also increases the overlap with conservation priorities identified for many other tropical plant and animal species (e.g., (*7*)). They are essential for maintaining biodiversity and ecosystem services; they disperse seeds, consume and pollinate plants, and create canopy gaps and trails (*8*). Finally, habitats important to apes, which mostly comprise tropical forests, play a crucial role for global climate change mitigation due to their ability to extract carbon dioxide from the air, create clouds, humidify the air and release cooling chemicals (*9*).

The IUCN Red List recently estimated that only 2-13% of all primate species were threatened by road and rail construction, oil and gas drilling, and mining, whereas 76% and 60% were negatively impacted by agriculture and logging, and wood harvesting, respectively (*4*). Similarly, mining currently ranks only forth in the frequency of reported threats across African ape sites documented in the A.P.E.S. Wiki (*10*), 65 out of 180 sites, i.e., 36% of all sites for which threats have been documented; (*11*) and is preceded by hunting (89% of sites), logging (62%), and agricultural expansion (62%). However, given recent findings on the density of mining areas across Africa (*2*), these values might be a considerable underestimation of the real threat of mining to apes. This discrepancy may be due to the lack of data from mining sites (i.e., only two of the 180 African ape sites included in the A.P.E.S. Wiki are mining areas as of March 2023). In addition, mining companies that conduct Environmental Impact Assessments (EIAs) typically practice data embargoes which prohibit use of the data by second- or third parties (see also 2022 Nature Benchmarks). As a result, there are few published studies that scientifically assess the impacts of mining on wildlife populations (*12*).

The direct and indirect impacts of industrial mining (hereafter ‘mining’) are manifold (Figure 1). Mining areas are highly dynamic and impactful activities already start during the exploration phase. During this phase, high noise production, caused by extensive drilling and blasting can disturb the communication of species, such as primates (*13*) and result in functional loss of otherwise intact habitat (*14*). Physiological responses to noise pollution have also been documented in various other wildlife species and include among others, increased heart rate, damage to the auditory system and ultimately, a decrease in survival probability (*15*). Removal of vegetation may already be initiated during this phase where very distinct drilling lines can often be visible from satellite imagery (*16*). During the exploitation phase, digging, blasting, and the use of heavy machinery typically result in direct impact within the project’s development area in the form of habitat destruction, fragmentation, and degradation (Fig 1.). The release of pollutants, such as heavy metals and toxic chemicals can contaminate air, water sources, and soil, potentially causing health issues (*17*–*19*) and disrupting food chains. While studies on the effect of light pollution are still scarce and non-existent for apes, a recent meta-analysis found that exposure to artificial light at night induces strong responses for physiological measures, (e.g., reduced melatonin levels), longer daily activity, and life history traits (e.g., reduced reproductive success), also in diurnal species (*20*).

**Fig. 1.**
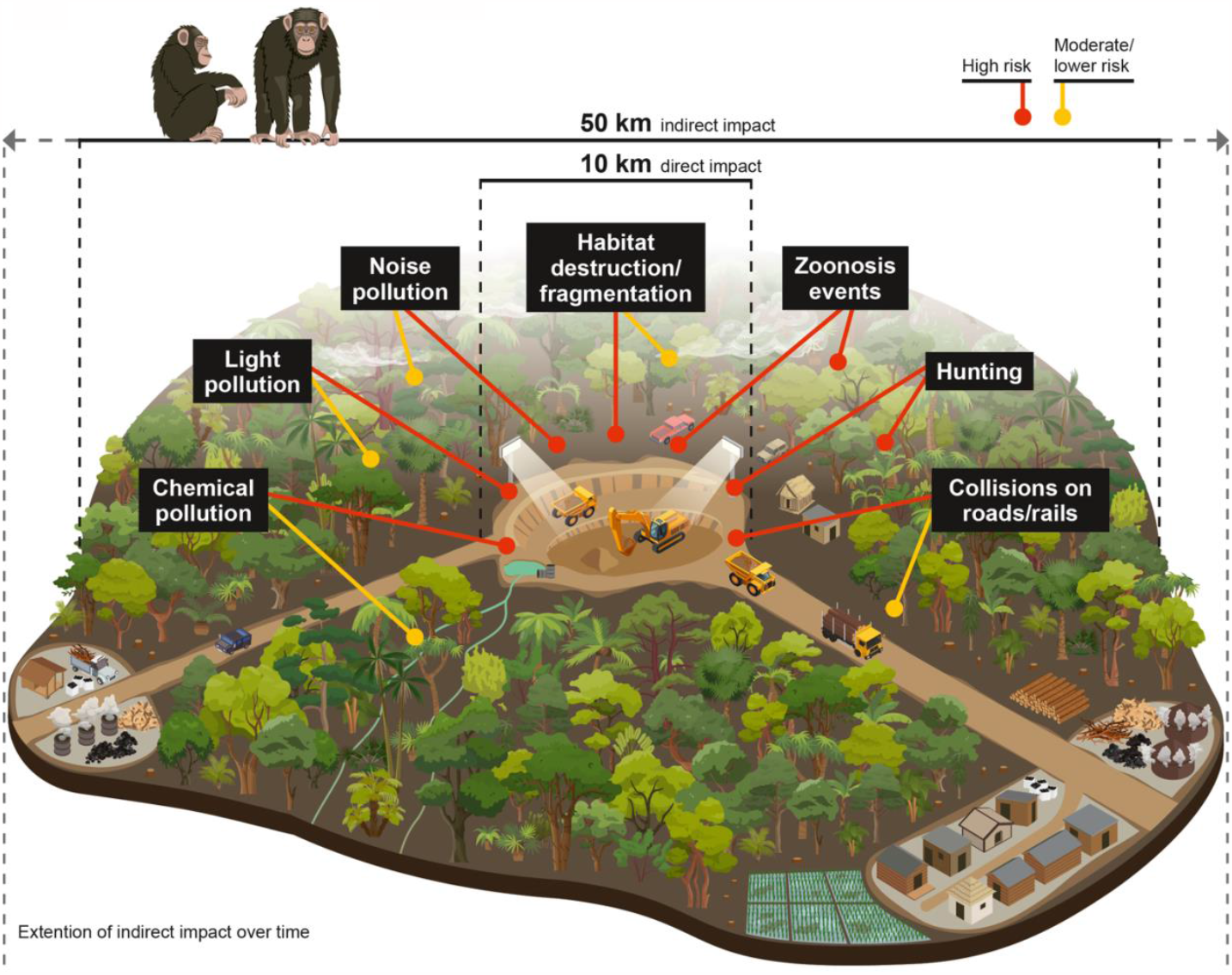
Schematic overview of the potential direct (10 km) and indirect threats (50 km) on apes linked to mining activities. Expected high and moderate to lower risk of impact is indicated by red and yellow pointers, respectively.

Indirect mining impact beyond the mining lease boundary is much more difficult to quantify and only a few studies on this topic have been published to date (e.g., (*21*–*23*), Figure 1). In 2017, Sonter and her colleagues demonstrated that large scale industrial mining operations caused significant deforestation over time and up to 70 km from mining lease boundaries in Brazil’s Amazon Forest. Furthermore, a recent global pan-tropical assessment found that in two-thirds of the 26 investigated countries, deforestation rates were higher close to the actual mining areas than in areas farther away, even when controlling for other known determinants of tropical deforestation (*24*). In some of these countries, the authors found high statistical significance for mining driving deforestation in the surrounding areas up to 50 km outside the mining areas. This is largely ascribed to in-migration of people and induced access resulting in an increased demand for land, charcoal, fuelwood, and roads (*23*).

Once extracted, many minerals are typically transported to the nearest port from where they are shipped to destinations around the world. Associated infrastructures, such as road and rail development therefore go hand-in-hand with activities in and around the concession site. The threat to wildlife posed by linear infrastructure is mostly indirect as demonstrated by numerous studies (e.g., (*25*–*28*)) however, collisions of vehicles with apes trying to cross the road have been reported previously (*29, 30*). Recently, Andrasi and colleagues (2021, (*31*)) estimated that western chimpanzee density is negatively impacted within a distance of about 16-19 km away from major-, and 5-6 km from minor roads. Various underlying threats negatively influence wildlife along roads: they include induced access, increased fire incidence, soil erosion, landslides, biological invasions, increased hunting pressure, and proliferation of agriculture (*32*). Finally, apes in mining areas are likely to have an increased risk of contracting disease from humans due to increased frequency in contact (*33*). This is aggravated by the fact that people and goods are moving more rapidly and further into remote locations potentially introducing diseases which were not known to those areas (*34*). However, an additional complex issue is the link between large scale development projects and the resulting habitat change and emergence and spread of diseases. Deforestation in tropical regions has often been associated with increased outbreaks of infectious diseases such as Dengue fever, malaria and yellow fever, some of these diseases affect great apes as well (*35*). The underlying mechanisms are often complicated: A study of zoonotic malaria, transmitted by macaques in Malaysian Borneo, confirmed the link between zoonotic spillovers and deforestation but showed complex and different effects of forest degradation at different scales (*36*).

To quantify the potential impact of industrial mining on wildlife population abundance, we used African great apes as a case study. They are particularly important in this context, since they are the only taxon specifically mentioned in the International Finance Corporation’s (IFC) Performance Standard 6 Guidance Note 73 (GN73) as a taxon that is likely to trigger so-called ‘Critical Habitat’ (CH), which imposes strict environmental regulations on mining companies that are seeking IFC funding (or loans from other lenders aligning with these standards) and that want to operate in these areas. It requires companies to reach out to the IUCN/ Species Survival Commission (SSC) Primate Specialist Group (PSG), Section on Great Apes for consultation (*37*). Specifically, mining projects operating in CH must implement mitigation measures to effectively counteract their ecological impact, ultimately resulting in a net increase in the overall population of great apes.

Using data spanning 17 Africa nations over an area of 1,507,811 km2, we estimated the magnitude of the potential direct and indirect negative impact from mining activities on ape abundance in and around operational and preoperational mining areas. To do this, we integrated a global mining dataset with range-wide estimates of ape density distribution. We investigated 1) how many African apes could potentially be negatively impacted by mining activities across their range, 2) whether mining areas often overlapped with high ape density areas, and 3) to what extent great ape survey data are available across these mining areas. Furthermore, we 4) quantified the spatial overlap of mining areas with likely Critical Habitat triggered by biodiversity features unrelated to apes and 5) identified hotspots of spatial overlap of high mining and ape densities.

## Results

### Geographical distribution of mining density in relation to ape density

High ape densities broadly coincide with operational and preoperational mining areas (with 50 km buffers) throughout most of the ape range in West Africa, in Gabon, southern and western Republic of Congo (from here on “Congo”) and southern Cameroon in central Africa, and in Uganda along the border of the Democratic Republic of Congo (DRC) (Fig. 2). Here, it is important to note that although artisanal mining poses a serious threat to apes and other wildlife in and around protected areas (e.g., Spira et al. 2019), it is not included in this analysis for reasons described in methods. Central Africa includes the largest percentage of areas with high ape densities outside of mining areas (63%), followed by East- (20%), and West Africa (14%), i.e., areas potentially not threatened by mining (Fig. S1). The most critical areas, i.e., those with relatively high ape densities (0.16- 6.07 apes/ km2, median= 0.3) and moderate to high mining densities (3- 42 mining areas/ km2; median= 3.8) are currently not protected (Fig. S2).

**Fig. 2.**
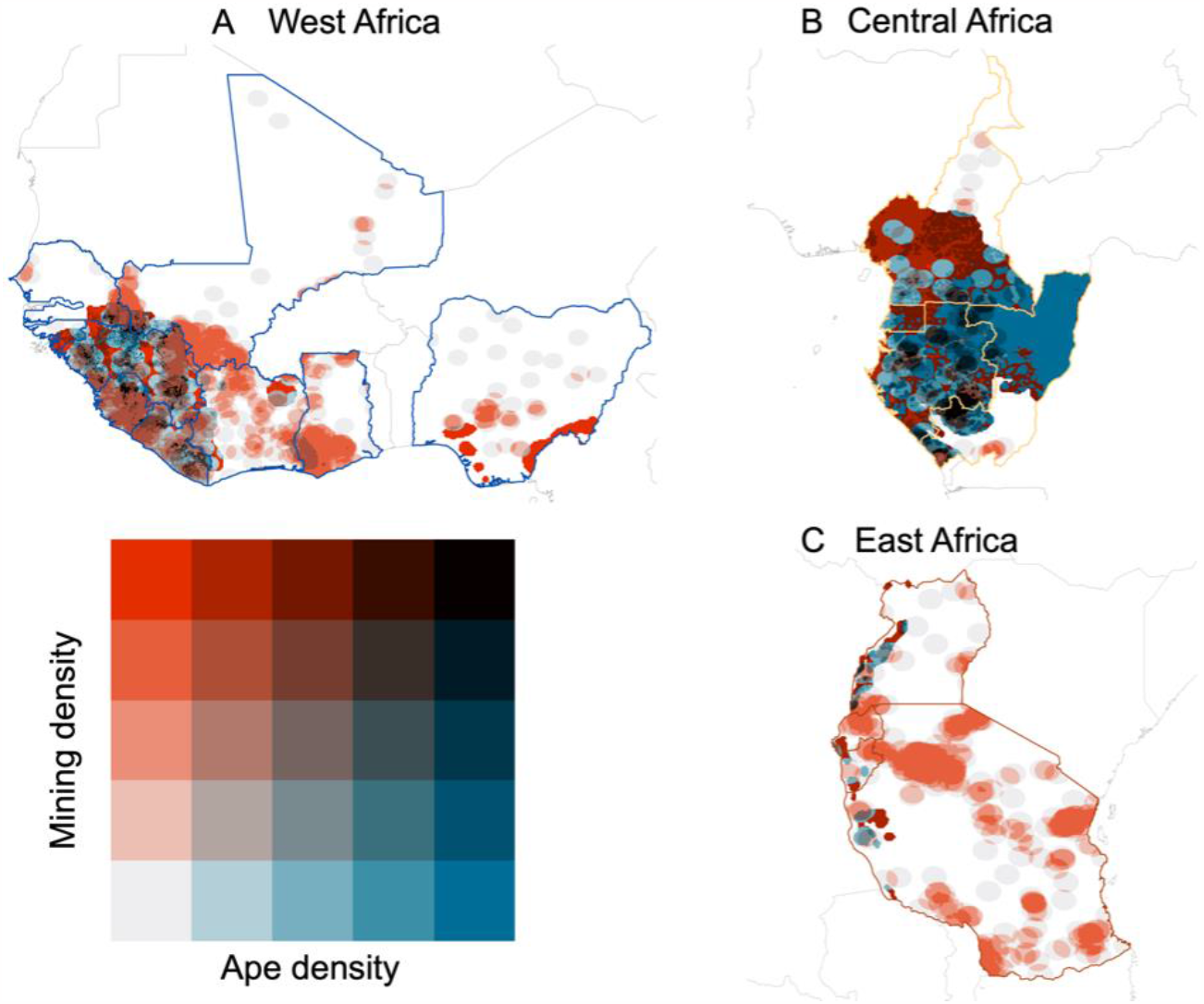
Bivariate choropleth showing the relationship between mining density (using 50 km buffers around mining sites) and ape density in (A) West Africa, in (B) central Africa, and in (C) East Africa. Each color change indicates a 20% quintile change in mining and ape density.

### Mining overlap with high-vs low ape density areas

Pre-operational-, and to a lesser degree, operational mining areas and their 10 km buffers in Liberia, Senegal and Sierra Leone in West Africa more often overlapped with high-rather than with low-ape density areas (Fig. 3). In these countries, chimpanzee range is either very restricted (i.e., Senegal) or chimpanzees are widely distributed but their range is highly fragmented (i.e, Liberia, Sierra Leone) and competition for different land uses is high. In countries with relatively large and/ or less fragmented ape populations, such as the Republic of Guinea (from here on “Guinea”) in West Africa and in Cameroon, Congo, Equatorial Guinea and Gabon in Central Africa, mining areas consistently had lower ape densities than non-mining areas. In Burundi and in Côte d’Ivoire, the majority of apes occur in a few protected areas, where industrial mining is less of a threat.

**Fig. 3.**
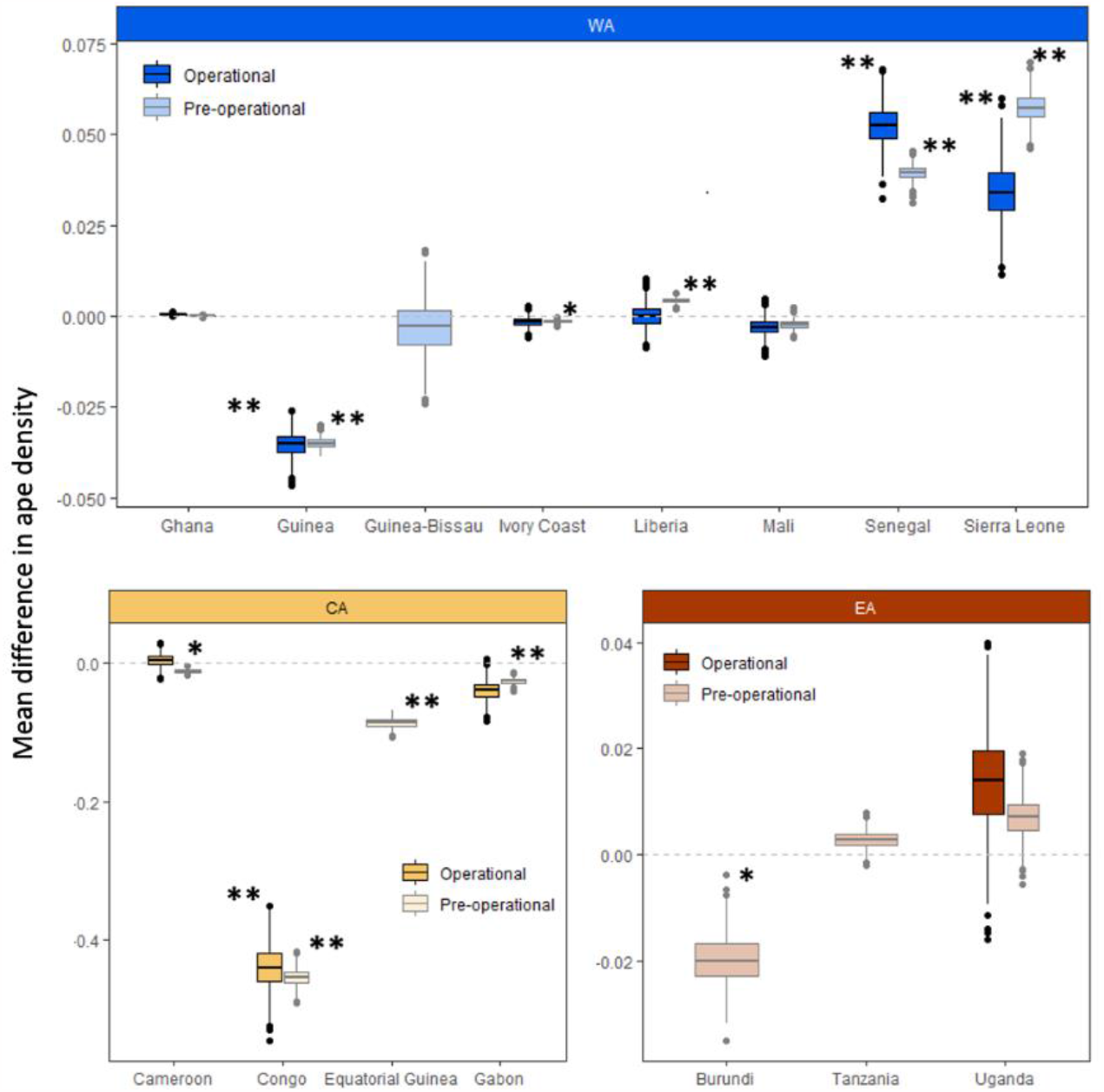
Box plots comparing the average difference in randomly selected samples of ape densities between areas within a 10 km buffer of preoperational and operational mining areas and randomly selected non-mining areas across countries in West Africa (WA), Central Africa (CA), and East Africa (EA). The dotted line indicates no difference between these areas. Values above the dotted line indicate that mining areas are located more often in areas with high-rather than low ape densities and vice versa. Nigeria and Rwanda are excluded as they do not include pixels that occur inside the ape range. Significant differences are marked with an asterisk (*p<0.01, ^**^p<0.001).

Positive spatial correlations between mining and ape density (i.e., more mining areas located in high-rather than low ape density areas) were observed more frequently when analyzed for mining areas with 50 km buffers (Fig. S3). When using 50 km buffers to reflect potential negative indirect impact of mining activities (see e.g., (*24, 38, 39*)), mining areas in five out of eight West African range countries (Guinea, Guinea-Bissau, Liberia, Mali, Senegal) overlapped more often with high-rather than low ape density areas within each of those countries. Mining areas in Tanzania and Uganda in East Africa, and in Gabon and Cameroon in central Africa, also more often overlapped with high-rather than low ape densities. Some relatively small countries (Burundi, Rwanda, Equatorial Guinea) and those with very small and spatially restricted ape populations (Côte d’Ivoire, Nigeria), showed the reverse pattern (i.e., mining areas overlapped more often with low-rather than high ape densities) and in Congo, a country with a very large and widely distributed ape population, mining areas consistently had lower ape densities than non-mining areas. In Ghana, there was no difference between operational and preoperational mining and non-mining areas, neither for 10, nor 50 km buffers, probably because of the extremely small population size (≅25 chimpanzees, (*40*)) and restricted area of this ape population. For detailed statistics of the t-tests refer to Table S1.

### Overlap of ape populations with mining areas

Mining areas and their 10 and 50 km buffers overlapped with 3% and 34% of the total ape population in Africa, respectively (Table 1). The spatial overlap of preoperational and operational mining areas with spaces important to apes was highest in West Africa, followed by East and Central Africa (Fig. 4). However, it is important to note that most of these areas (84.6%) represent mineral exploration areas (i.e., pre-operational mining areas), which may or may not become operational in the future. Countries with the largest overall overlaps in ape population abundance and mining areas (in terms of numbers of apes potentially affected) included Gabon, Congo and Cameroon in Central Africa, and Guinea in West Africa (Table S2). Although our dataset included fewer mining areas in Central-(12% of total mining areas) than in East-(27%) and West African range countries (61%), more individual apes would potentially be threatened by mining in this region, because of higher overall ape densities in this region (*41*). Countries that had the largest proportional overlaps between ape population abundance and mining areas (in terms of proportion of population potentially affected), included Liberia, Sierra Leone, Mali, and Guinea, are all located in West Africa (Fig. 4). Therefore, Guinea had one of the largest proportional- and overall overlaps of mining- and chimpanzee density, where >23,000 individuals or up to 83% of Guinea’s population could be directly or indirectly influenced by mining activities soon. All country-specific overlap statistics are available in Table S2.

**Table 1.** Total- and proportional overlap between ape density distribution and mining areas with 10 km and 50 km buffers in West-, central-, and East Africa.

| Region | No. apes potentially threatened by mining (10 km buffers) | Proportional overlap (10 km buffers) | No. apes potentially threatened by mining (50 km buffers) | Proportional overlap (50 km buffers) |
| --- | --- | --- | --- | --- |
| West Africa | 5,691 | 12% | 39,599 | 82% |
| Central Africa | 10,711 | 2% | 135,042 | 29% |
| East Africa | 292 | 4% | 4,175 | 62% |
| **Total** | **16,694** | **3%** | **178,816** | **34%** |

**Fig. 4.**
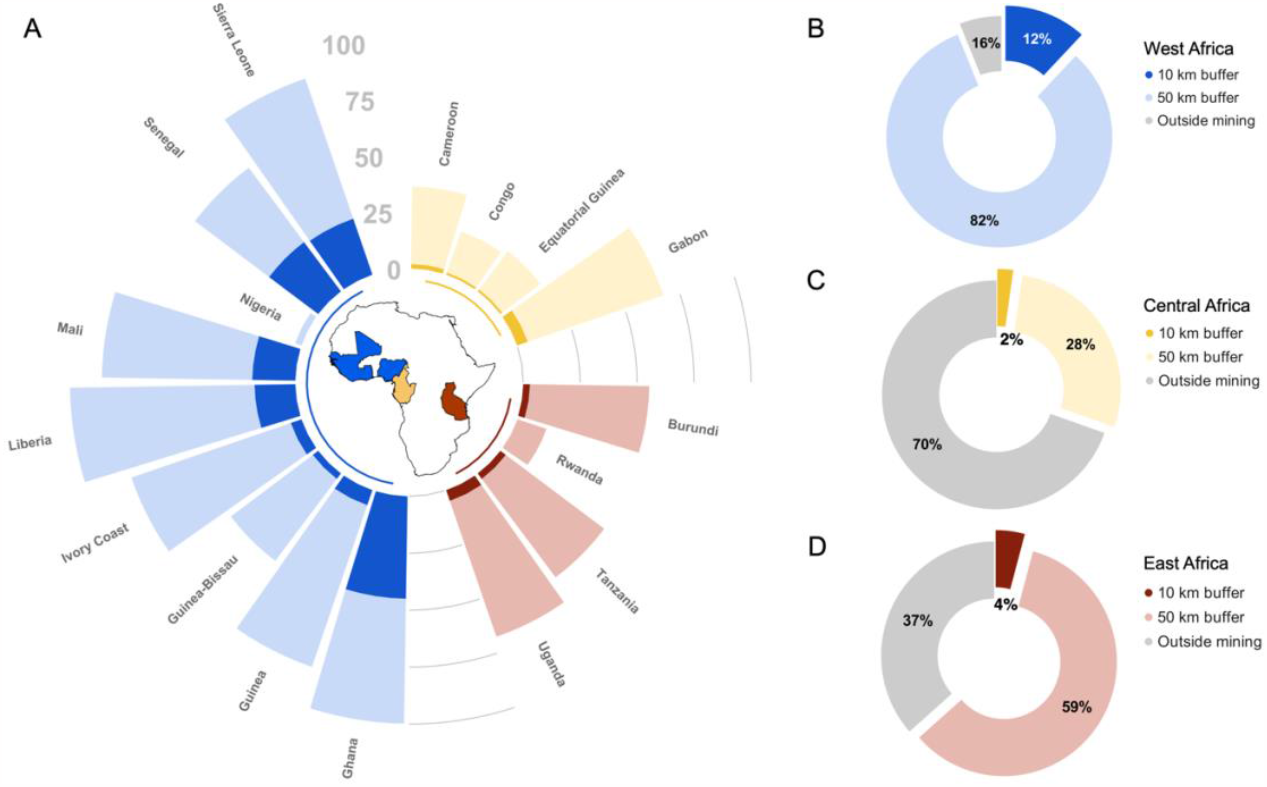
(A) Proportion of ape population threatened by mining (operational and preoperational mining areas) with a 10 km buffer (dark shades) and with a 50 km buffer (light shades) for range countries in the different regions. Total regional estimates of the proportion of ape populations threatened by mining in **(B)** West Africa, in **(C)** central Africa, and in **(D)** East Africa.

### Overlap with Critical Habitat triggered by biodiversity features other than apes

We found that 20% of mining areas with 10 km buffers overlapped with potentially additional Critical Habitat (CH) triggered by biodiversity features other than apes (Fig. S4). When we compared CH- to ape density distribution, we found large areas that did not overlap between these two layers (Fig. S5). This discrepancy is most profound in Guinea and Sierra Leone in West Africa, and in Congo and Gabon in central Africa.

### Availability of ape data for mining areas

At the time of analysis, only 3% of pixels included in mining areas had survey data stored in IUCN SSC Ape Populations, Environments and Surveys (A.P.E.S.) database (*10*) and only 1% of the total area surveyed and archived in the A.P.E.S database overlapped with operational or preoperational mining areas (Fig. S6).

## Discussion

Corporations and their operations are the most important contributors to world-wide biodiversity loss and ecosystem destruction (*42*). Mining is one of the top drivers of deforestation globally with tropical rainforests standing out as mining-induced deforestation hotspots (*24*). Moreover, deforestation within current mining leases suggests that the rate of mining-related forest loss has increased significantly over the past 10 years (*24*). These patterns, which are driven by a rapidly growing global demand for critical metals vital to energy transitions, are expected to exacerbate deforestation over the coming years if companies continue business-as-usual. Until now, private sector contributions to a more sustainable and nature-positive future have remained low. In a recent ranking published by the World Benchmarking Alliance (*43*), only 5% of the 400 assessed companies had carried out science-based nature and biodiversity impact assessments of their operations and business models.

To address these issues, the Sustainable Critical Minerals Alliance (SCMA) was announced at COP 15. Its work plan, funded by member countries and private sector partners, focuses on four key areas: 1) promoting responsible mining practices, 2) developing new low-impact technologies, 3) creating circular economies for critical minerals, and 4) sharing benefits equitably. Related to key area 1, this study provides species-level data on the potential threat of mining on population abundance across the entire range of African great apes, a taxon threatened by extinction and of high ecological, economical and anthropological significance. Our results indicate that the magnitude and extent of the potential threats of mining on apes in Africa has been grossly underestimated. In many instances and throughout their range in Africa, preoperational and operational mining areas coincide with areas of high importance to apes, where many of these overlapping areas currently lack adequate protection measures (Fig. 2). Although DRC was not included in our analyses, there is evidence that mining has had significant impacts on the Eastern chimpanzee (*Pan troglodytes schweinfurthii*) and Grauer’s gorilla (*Gorilla beringei graueri*) population inside and outside protected areas, supporting our results (*44*).

This pattern was particularly profound in West Africa, which was the region with the largest number of mining areas. Here, ape range is highly fragmented and spatially restricted and areas with large mineral deposits that are not yet developed, are directly competing with areas that are important to apes. Furthermore, great ape densities were significantly higher inside than outside mining areas and their 10 km buffers in three out of eight West African range countries (Fig. 3), and in five out of eight countries when using 50 km buffers (Fig. S3). We estimated that more than one-third of the entire great ape population in Africa - nearly 180,000 individuals - could be directly or indirectly threatened by mining now and in the near future. Apes in West Africa could be most severely affected, where up to 82% of the population currently overlaps with operational and preoperational mining areas and their 50 km buffers (Fig. 4).

Given the increase in overlap between areas developed by mining projects and areas preserved in their natural state to protect apes and other threatened wildlife species, we have to substantially step up our efforts to integrate conservation goals with economic development targets. The ‘mitigation hierarchy’ (*37, 45*), as articulated by the Business and Biodiversity Offsets Programme (BBOP) and the IFC, is a best practice approach to managing potential impact on biodiversity by development projects that receive funding from IFC or other lenders that align with their standards. This approach advocates applying efforts early in the development process to avoid adverse impacts to biodiversity wherever possible, then reduce impacts that cannot be avoided, rehabilitate impacted areas; and then compensate for any residual impacts (*46, 47*). However, mining companies frequently only apply measures to mitigate a) direct impact b) during exploitation and c) within the mining lease boundaries. They fail to consider that their impacts, whether direct or indirect, occur during all project development stages and spill over to a wider geographic area. To allow ape populations to disperse and relocate, mitigation of both direct and indirect impact should extend beyond the administrative boundaries of the mining project. At the same time, companies should make a greater effort to identify and anticipate indirect impacts induced by e.g., mining-related human in-migration and zoonotic disease transmission. The time frame over which a net gain in ape population abundance is achieved is also all too often underestimated. If the time frame is too short, the populations may not have enough time to increase sufficiently.

Considering the complex social organization and dynamics of African great apes and associated elevated risks of mortality, as well as the paucity of suitable release sites, translocation of groups from highly impacted areas is not a feasible option (*48*). Additionally, translocation and relocation of wildlife potentially raises several ethical and legal issues due to the stress inflicted by the animal and risks associated with starvation and predation by other species (*49*). Finally, restoring habitat simply takes too long for resident apes to benefit from this intervention. Therefore, unless great ape habitat is avoided entirely, mitigation is unlikely to prevent ape population declines. Companies should therefore reconsider the long-term feasibility of exploration sites in areas important for apes, due to their environmental responsibilities and the costs associated with achieving no net loss/ net gain in ape abundance. Also, lending banks should refrain from funding projects in these areas. To illustrate this, if corporations ceased their exploratory activities in areas likely to contain a minimum of 20 apes, this would result in 38% (22 out of 58) of putative mining projects situated within the African ape habitat to remain undeveloped. Notably, nine of these areas exhibit the potential to accommodate populations exceeding 50 apes.

To compensate for any residual impact that could not be avoided, reduced, or restored, mining companies can implement compensation measures by creating biodiversity-offsets to ensure that an equal or greater area of identical habitat or ape population is protected or improved (*50*). However, offsets are controversial and their effectiveness for apes has yet to be demonstrated (*51*–*53*). Offset design and implementation is frequently guided by company internal standards, lending banks, or international best-practices and few African ape-range countries have national policies guiding or requiring offsets (*51*). A recent independent assessment by the ARRC Task Force of the Section on Great Apes and Section on Small Apes of the IUCN SSC Primate Specialist Group (*54*) has shown that even the most ambitious and cutting-edge efforts by the private sector to offset residual impacts on apes and their habitat are not sufficient to effectively mitigate the total loss they incur to great ape populations. One key factor is the duration of offsets, which is often set equal to the length of exploitation activities (generally c.20 years). This time period is too short to achieve any significant gains for apes. These temporary actions do not ensure long-term conservation of apes, while most impacts at the mining sites are permanent. Offsets also do not consider impacts from mining exploration activities, and legacy impacts when projects are sold to different companies.

Where compensation schemes are considered, offsets must be designed in such a way as to take into account the cumulative threats across the landscape or region, ideally forming part of existing national or regional conservation strategies. The estimates provided in this study could serve as a first approximation based on which an initial screening for suitable aggregated offset schemes could be conducted. Our study also provides some guidance with regards to where to compensate for residual impact. Investing in increased protections might be more feasible where high ape densities exist outside of mining sites. Alternatively, aggregated offset strategies could focus on contributing to existing protected areas to improve their effectiveness (e.g., by financially investing in management activities and staff) (Fig. S1).

We also found that 20% of mining areas overlapped with areas that likely qualify as Critical Habitat triggered by biodiversity features other than apes (Fig. S4), which, according to international regulatory frameworks, would hinder projects from receiving financial support (i.e., (*37*). Similarly, another study found that 32% of all mammal species worldwide with more than 30% of habitat within mining areas are currently listed as Threatened with extinction on the IUCN Red list (*55*). Since species of conservation concern would likely trigger CH status, companies operating in these areas should have adequate mitigation and compensation schemes in place to minimize their impact, which seems unlikely, given that most companies seem to lack robust species baseline data (*43*). What is of even greater concern is the spatial overlap between areas set aside for conservation and those potentially influenced by mining. For example, it is estimated that 8% of the global area potentially influenced by mining overlaps with protected areas, 16% with Remaining Wilderness and 7% with Key Biodiversity Areas (*2*). Another study that examined the intersection of mines with protected areas identified 2558 boundary violations totaling about 6,232 km^2^, or 9.5% of all areas identified as mining projects (*56*). This is supported by the information on world-wide downgrading, downsizing, and degazettement of protected areas (PADDD), providing evidence for more than 3,000 enacted cases of PADDD in nearly 70 countries, covering about 1,300,000 km^2^ (updated from (*57*)).

Our results confirmed the lack of data sharing by mining projects, where only 1% of the ape survey data from Africa that is currently stored in the A.P.E.S database – the first and only public repository for data from surveys of apes and their habitats – was collected in and around mining areas (Fig. S6). This lack of transparent data sharing hampers science-based quantification of impacts of mining on apes and their habitat and the development of effective mitigation strategies. This was reflected in the results of the first global synopsis of the effects of primate conservation interventions examining approximately 13,000 publications, which found a marked absence of studies on the effectiveness of conservation strategies specifically designed to reduce the impact of mining on apes (*58*). We therefore stress the need for mining companies to make their biodiversity data publicly available in a central database, such as IUCN SSC A.P.E.S. or the Global Biodiversity Information Facility (GBIF) and call on the International Finance Corporation (IFC) and other regulatory frameworks to urge companies to provide access to their data.

The large overlap between mining areas and areas important to apes is partly because many of the minerals needed for the energy transition are in places that have not yet been industrialized, which typically include rural or remote parts of the world. This means that current climate solutions could lead to more industrialization in these places, which could worsen the climate crisis (*59*). The production of biofuels from food and feed crops exemplifies this paradox, where increases in bioenergy cropland to meet global demands in biofuel are expected to cause severe impacts on biodiversity that are not compensated by lower climate change impacts (*60*). In addition, the injustices inflicted by the expansion of industrial development are already immense (*61*) and may worsen with an increase in unsustainable economic development in previously undeveloped areas (*62*). To illustrate this, 69% of energy transition minerals and metals projects worldwide are on or near land that belongs to Indigenous people or small holder farmers and pastoralists, with an even higher proportion (77%) of overlap in Africa (*59*).

The Sustainable Critical Minerals Alliance (SCMA) is a significant step forward in the global effort to ensure that the transition to a low-carbon economy is sustainable and equitable. However, Africa’s great apes and many other threatened wildlife species are at high risk from industrial mining activities, which are likely to increase as the world transitions to a low-carbon economy. The inclusion of great apes in IFC’s Performance Standard 6 Guidance Note 73 and the creation of the ARRC Task Force, comprised of foremost experts in ape conservation tasked with offering independent guidance on how to mitigate the adverse effects of energy, extractive, and related infrastructure projects on apes, instills optimism that efforts to integrate conservation goals with economic development targets are increasingly being taken up by environmental policy and private investors. Our findings highlight the need for the mining sector to increase transparency and make their environmental data more accessible. This would allow for better independent assessments of the risks posed by mining activities to endangered flora and fauna. We also call upon companies, lenders, and nations to reevaluate investments in exploration activities in areas of high biodiversity and recognize the greater value of leaving some regions untouched by industrial activity, as these actions are vital for preserving ecosystem services, preventing disease spillovers addressing future epidemics or pandemics, and mitigating climate change.

### Limitations of the Approach

A limitation of this study is that the mining data set that we used did not include artisanal mining areas. Although small-scale, informal and artisanal mining areas constitute only 1.63% of the total mine area globally, the proportional magnitude of the artisanal mining footprint is likely substantial, because these areas are often associated with severe environmental risks and no ecological protection measures (*56*). Therefore, our estimates of the impact of mining activities on apes in operational mining areas might be an underestimate of the true impact. Adding to this, mining activities have been observed to cause indirect impacts that expand across space and persist over time, as evidenced by a study conducted by Tang and Werner in 2023 (*56*).

On the other hand, the majority of mining areas included in this study are still in the exploration phase (proportion preoperational mines: West Africa= 81.6%, central Africa= 91,7%, East Africa= 87,8%) and it can be expected that not all of the preoperational mining areas will become operational in the future. A number of studies estimated the success rate of mining exploration (i.e., the proportion of exploration sites that become extraction sites) and calculated that the likelihood of discovery of a major deposit in areas where little to no previous mining activity has occurred, ranges from 0.3 - 0.5%, and is 5% in areas where mining activities have taken place previously (*63*). However, the geological potential of a site is not the only factor determining the success of a mine and other aspects, such as economic viability, market demand, social acceptance, global economic conditions, and regulatory and environmental factors, among others, influence return on investments in mining. While the return on investments is less than 1% globally, for Australia and Africa returns on investments in mining are considerably higher and estimated at 12% and 38%, respectively (*63*). Also, while a mine might be regarded as economically unfeasible at one point in time, it may become feasible at another point in time (e.g., as demand or the price for the mineral increases).

Likewise, operational mines may be implementing effective mitigation measures, thereby not impacting all great apes within 10 or 50 km buffers. Since the effects of these processes are difficult to quantify with the limited data at hand, they were also not considered in this analysis. Furthermore, since robust data on the extent of the direct and indirect impact of mining activities on apes in Africa are lacking, the buffers used in this study are mere approximations of true impact. In some instances, they may be an overestimate, e.g., in the case of relatively recent mines, or mines that have implemented appropriate avoidance and minimization measures, and in other cases they may underestimate the true impact of the mine, e.g., well-established and relatively large-scale operational mines. Finally, another source of uncertainty is the highly dynamic nature of impact from mines. A mine life cycle may involve periods of expansion followed by periods of reclamation or revegetation, further complicating the interpretation of results. Despite these limitations, we think that the results presented in this study provide a useful first global assessment of the potential threats of mining on apes in Africa.

## Materials and Methods

### Study Design

We used various data sources for analysis related to mining- and ape density in different geographical locations (Table 2). We used Mollweide equal area projection to analyze all data listed in Table 1 and matched all spatial layers at a 1*1 km pixel resolution. We combined great ape density distributions modeled by (*40*) and (*7, 41*), and mapped this for each ape range country in Africa (referred to as “range country” throughout the text). We excluded the DRC and the Central African Republic from analysis due to a lack of ape density information (*41*). However, in DRC there is extensive mining occurring within the Eastern chimpanzee and Grauer gorilla’s range, inside and outside protected areas, and thus the impact on their population is likely significant (*44*).The metric for ape density distribution is the number of apes per pixel.

**Table 2.**
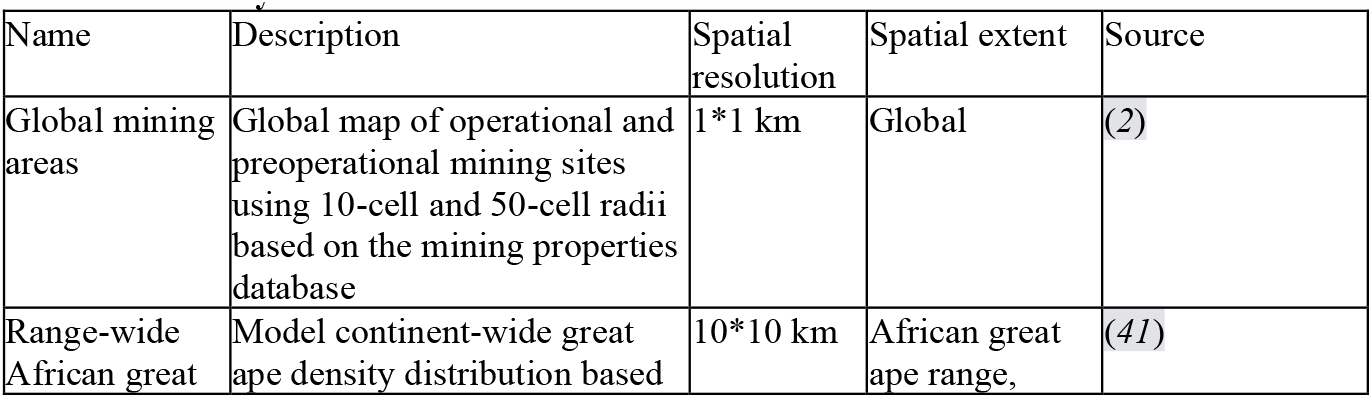
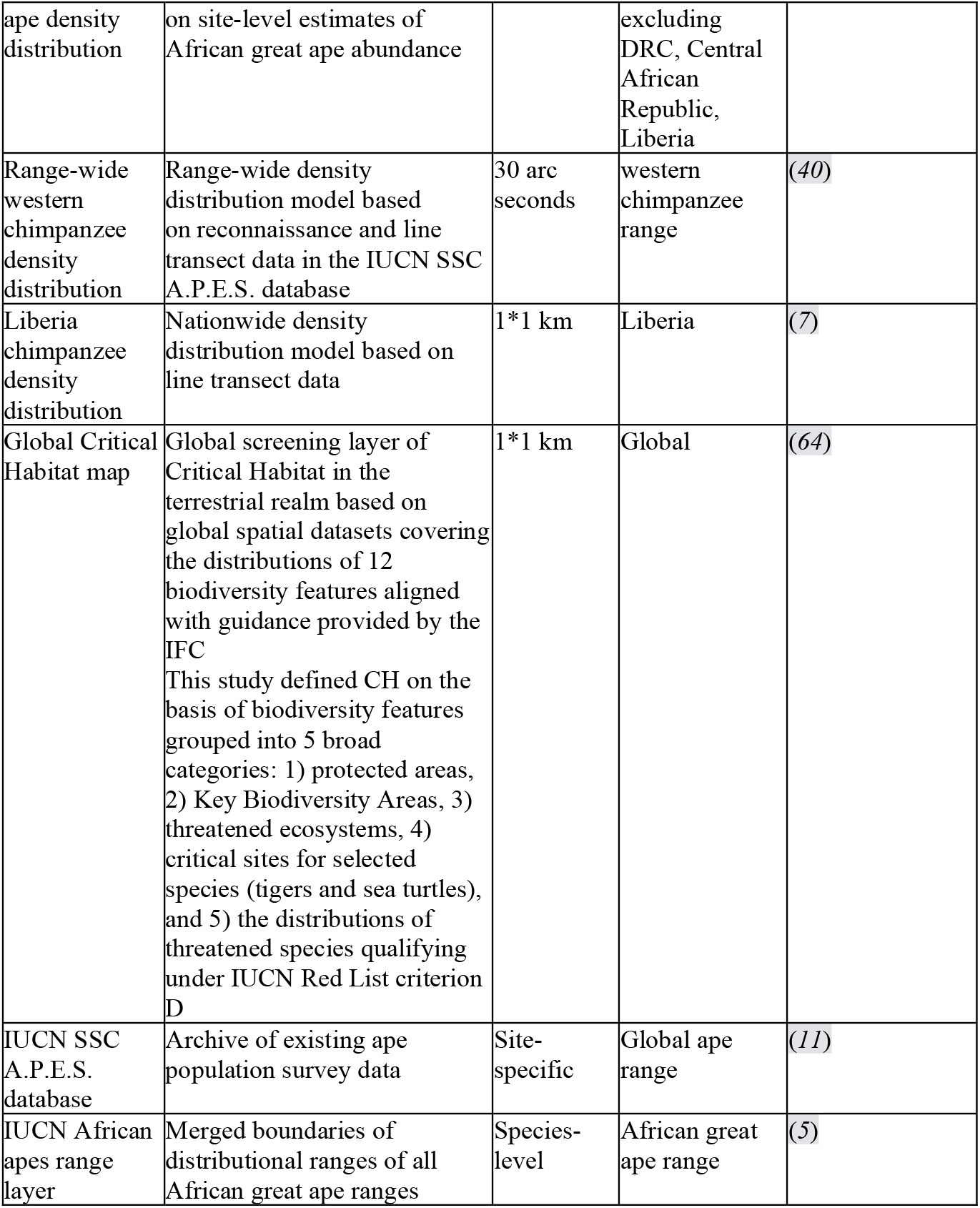
Name, description, spatial resolution, spatial extent, and source of datasets used in this analysis.

We had two mining datasets: a dataset that included industrial 1) preoperational (i.e., exploration) and 2) operational (i.e., exploitation) mining sites both with a 10-cell and a 50-cell radius, collectively referred to as “mining areas” throughout the text (*2*). Values in these spatial layers estimate mining density (i.e., number of overlapping mining sites per pixel). Since none of the preoperational sites are currently being mined, we use these as a proxy for potential future mining sites, recognizing that some of these sites may never be developed. We converted mining densities to binary values to indicate mining influenced areas where mining density was > 0.

### Buffer Areas

The global dataset on mining sites used in this study includes point locations only, and as such, the boundaries of the mining concession were not known. We therefore defined buffers to reflect the approximate extent of direct and indirect impacts from mines. To do this, we considered the results of previous studies that estimated average mining area, which is the area likely to be included within mining lease boundaries and which ranges from 0.36 to 12.3 km2. The study that estimated average mining area sizes at > 12.3 km2 focused on larger-scale operations (*65*), whereas studies that reported average mining area sizes at < 2 km2 included artisanal mining areas (*16, 56, 66*–*68*). Since the dataset used in this study provides coordinates of larger scale mines, and because mining-related threats like light and noise pollution or hunting cannot be visualized from satellite images, we believe that using a 10 km buffer to approximate the direct impact from mines is justified. Indirect impact of mining, on the other hand, has commonly been assumed to extend 50-70 km beyond the boundaries of mining areas (*24, 38, 39*) and we therefore decided to use a 50 km buffer to assess potential indirect impact of mining on African great apes.

### Statistical Analyses

Geographical Distribution of Mining Density in Relation to Ape Density We used mining density, i.e, the number of mining areas (i.e., point locations and their 10 and 50 km buffers) overlapping with each pixel, and mapped this in relation to ape density distribution across range countries. Here, we merged operational and preoperational mining areas. We then grouped the values for mining density and ape density into quintiles and classified each pixel depending on the product of these two factors, resulting in a total of 25 classes. We mapped these 25 classes over geographical space to visualize pristine ape habitat (low mining density; high ape density) vs. ape habitat threatened by mining (high mining density; high ape density) vs. areas where mining does not threaten ape populations (high mining; low ape densities) and where neither mining-, nor ape densities are high. We excluded areas with values of zero for both mining and ape density. For each region, we also plotted the percentage area across quintiles of varying ape- and mining density, where we restricted this analysis to areas with high ape-densities (i.e., including only ape density values that fell into the fifth quintile) and across all five quintiles of varying mining density.

### Mining Overlap with High-vs Low Ape Density Areas

To compare ape density differences between mining and non-mining areas across the ape range and within each country, we first overlaid mining areas with ape densities. We then compared the distribution of ape densities from pixels that overlapped with mining areas to those outside of mining areas, but within the ape range. To account for the large variation in pixel numbers between countries, as well as mining and non-mining areas (mining areas always had much fewer pixels than non-mining areas), we selected the total number of pixels from within mining areas and randomly selected the same number of pixels without replacement from non-mining areas separately for each country. We then performed a t-test, repeating the process for 1,000 iterations, to determine if there were density differences between mining and non-mining areas. The large number of iterations and random selection approach minimized the likelihood of biased results stemming from specific pixel selection and resulted in more representative samples. This process was done separately for pre-operational and operational mines and for mining areas with 10 and 50 km buffers.

### Overlap of Ape Populations with Mining Areas

We overlaid the mining areas with ape density distribution and summed the number of apes estimated for each pixel at 1*1 km resolution to estimate the proportion of total ape population potentially threatened by current and future mining activities in each region and range country. Each pixel in the ape density distribution layer was weighted by the amount of overlap with mining areas. If, for example, 30% of the pixel area fell into a mining area, then only 30% of the number of apes in that pixel was included in the overall estimate of threatened apes per region and range country.

### Overlap with Critical Habitat Triggered by Biodiversity Features Other than Apes

We followed the procedure described in section 3.3.3 and summed the number pixels at 1*1 km resolution to estimate the proportion of area identified as likely and potential CH triggered by biodiversity features other than apes (Global Critical Habitat map; Table 1), that overlapped with operational and proposed mining areas in each region (West Africa, East Africa, central Africa) and range country. Each pixel in the Global Critical Habitat map was then weighted by the amount of overlap with mining areas. To investigate how likely CH triggered by the occurrence of apes complemented (or not) the areas identified as likely or potential CH triggered by biodiversity features other than apes, we compared the Global Critical Habitat map (clipped to range countries) with ape density distribution. This allowed us to identify additional areas of likely CH not yet included in the output maps produced by Brauneder and colleagues (2018, (*64*)).

### Availability of Ape Data for Mining Areas

We consulted the data in the IUCN SSC A.P.E.S. database (hereafter referred to as ‘A.P.E.S.’; (*11*)) to determine whether survey data existed for sites that overlapped with mining areas and their 10 km buffers. Here, we only included mining areas within the distributional range of great apes ((*5*), Table 1). To know if an ape survey was conducted in the area or not (which also included surveys that did not report the presence of apes in the area), we mapped all observations recorded during surveys over the global mining areas layer (Table 1) and calculated the proportion of pixels included in mining areas for which survey data were available (i.e., via request to A.P.E.S.). Here, we also included in the analysis the DRC and the Central African Republic since we assessed the spatial overlap of survey data from A.P.E.S. (and not ape densities as in the previous analysis) with mining areas.

### Data Processing

All analyses were performed in R (Version 4.2.0) using the following R packages: ‘raster’ (*69*), ‘terra’ (*70*), ‘sp’ (*71*), ‘sf’ (*72*), ‘rgdal’ (*73*), ‘ggplot2’ (*74*)), and ‘dplyr’ ((*75*), ‘tidyr’ (*74*), and ‘reshape2’ (*76*). In addition, we used QGIS (V 3.26.2) and ArcMap (V 10.7.1) to spatially visualize our data on maps.

## Acknowledgments

We would like to thank Emily Wendt for helping with formatting the manuscript and Gabriele Rada/ iDiv for helping with graphic design. We are also grateful to Ekwoge Abwe for comments on the first draft of this manuscript.

## Author contributions

Conceptualization: JJ

Methodology: JJ, LQ, JV

Investigation: JJ, LQ, JV, TS

Visualization: JJ, LQ, JV

Writing—original draft: JJ

Writing—review & editing: JJ, LQ, JV, MA, AB, GC, SH, TH, CYK, HK, IO-N, HMP, HR, JR, LS, TS

## Data and materials availability

All data will be made available in the main text or the supplementary materials upon acceptance of the manuscript.

