## Supplementary Text; Figs. S1 to S6; Tables S1 and S2 for "Threat of mining to African great apes"

**This PDF file includes:**

Supplementary Text
Figs. S1 to S6
Tables S1 to S2

**Supplementary Text**

Additional figures and tables including extended results.

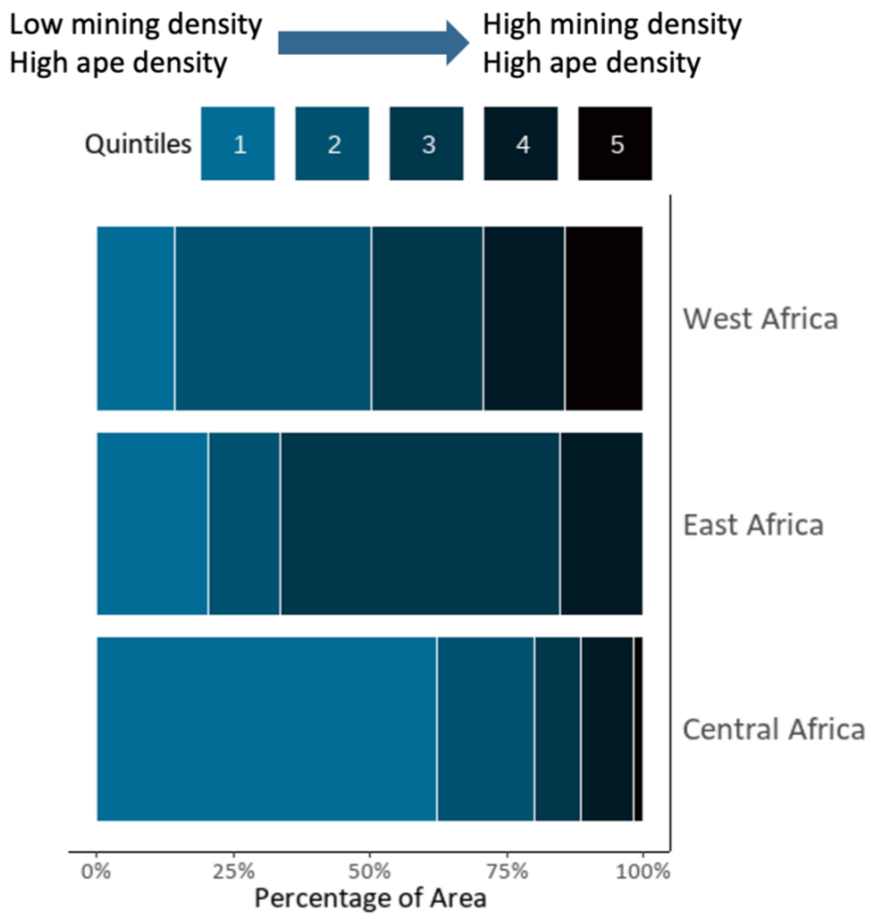


**Fig. S1.**

Percentage of area with high ape density that falls into quintiles with varying mining density. Each quintile represents a 20% change in mining density.

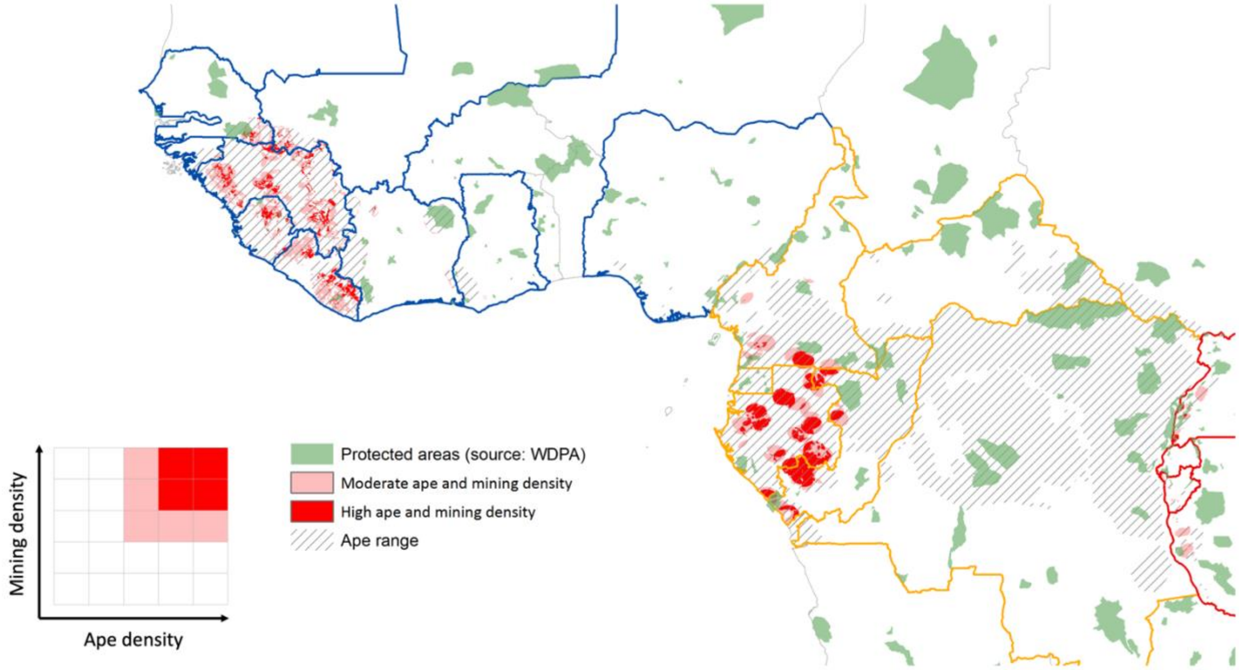


**Fig. S2.**

Distribution of so-called critical areas (dark red), which are areas with high ape and high mining density in relation to the location of IUCN protected areas (categories 1-4). Categories are based on mining and ape density quintiles.

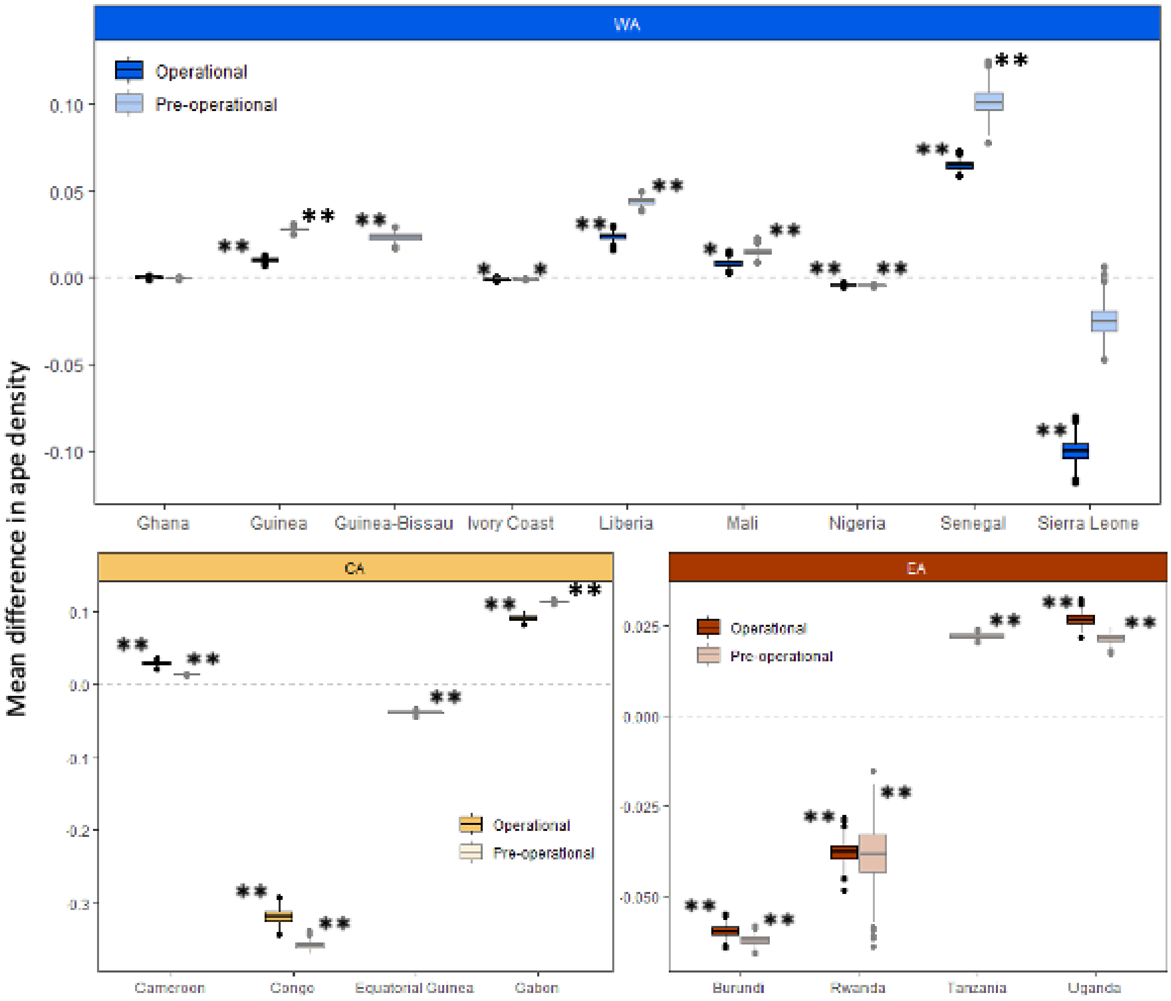


**Fig. S3.**

Box plots comparing the average difference in randomly selected samples of ape densities between areas within a 50 km buffer of preoperational and operational mining areas and randomly selected non-mining areas across countries in West Africa ("WA"), central Africa ("CA"), and East Africa ("EA"). The dotted line indicates no difference between these areas. Values above the dotted line indicate that mining areas are located more often in areas with high- rather than low ape densities and vice versa. Significant differences are marked with an asterisk (*$p<0.01$, **$p<0.001$).

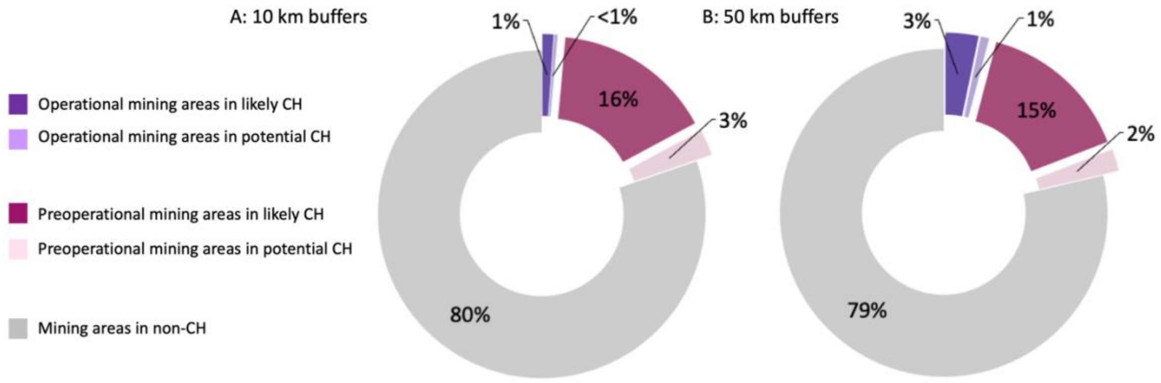


**Fig. S4.**

Proportional overlap of likely and potential CH with pre-operational and operational mining areas with (A) 10 km and (B) 50 km buffers.

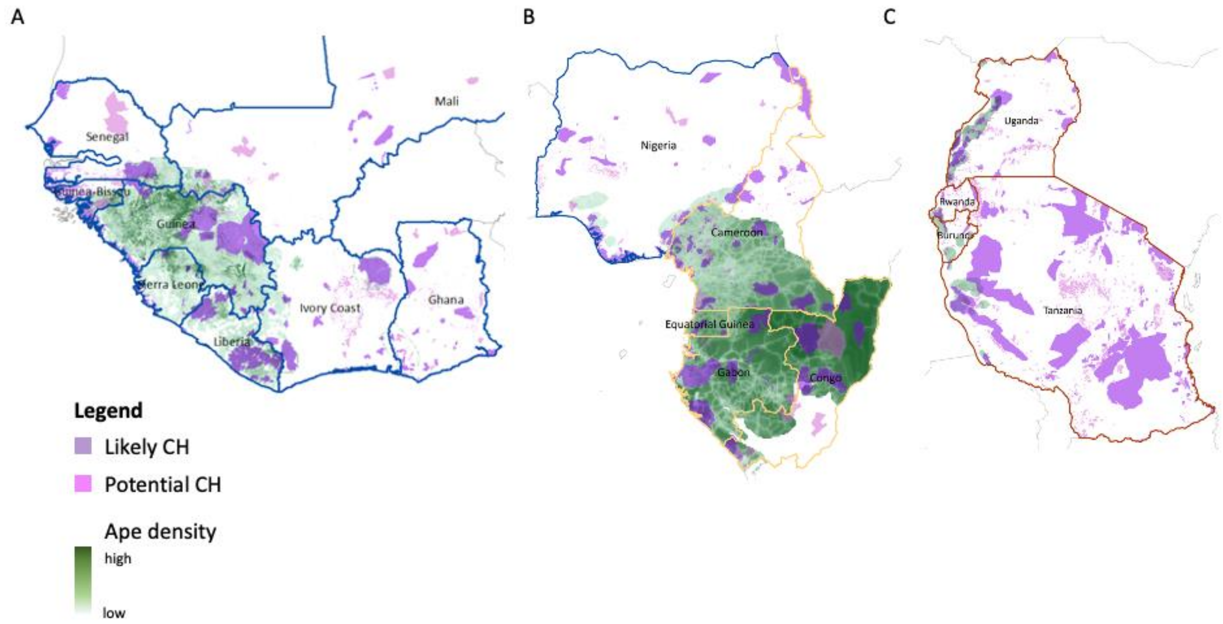


**Fig. S5.**

Distribution of likely and potential Critical Habitat (CH) mapped by Brauneder et al. (2018) in relation to chimpanzee density distribution in (A) West African range countries, (B) central African ape range countries and in Nigeria in West Africa, and (C) in East African ape range countries.

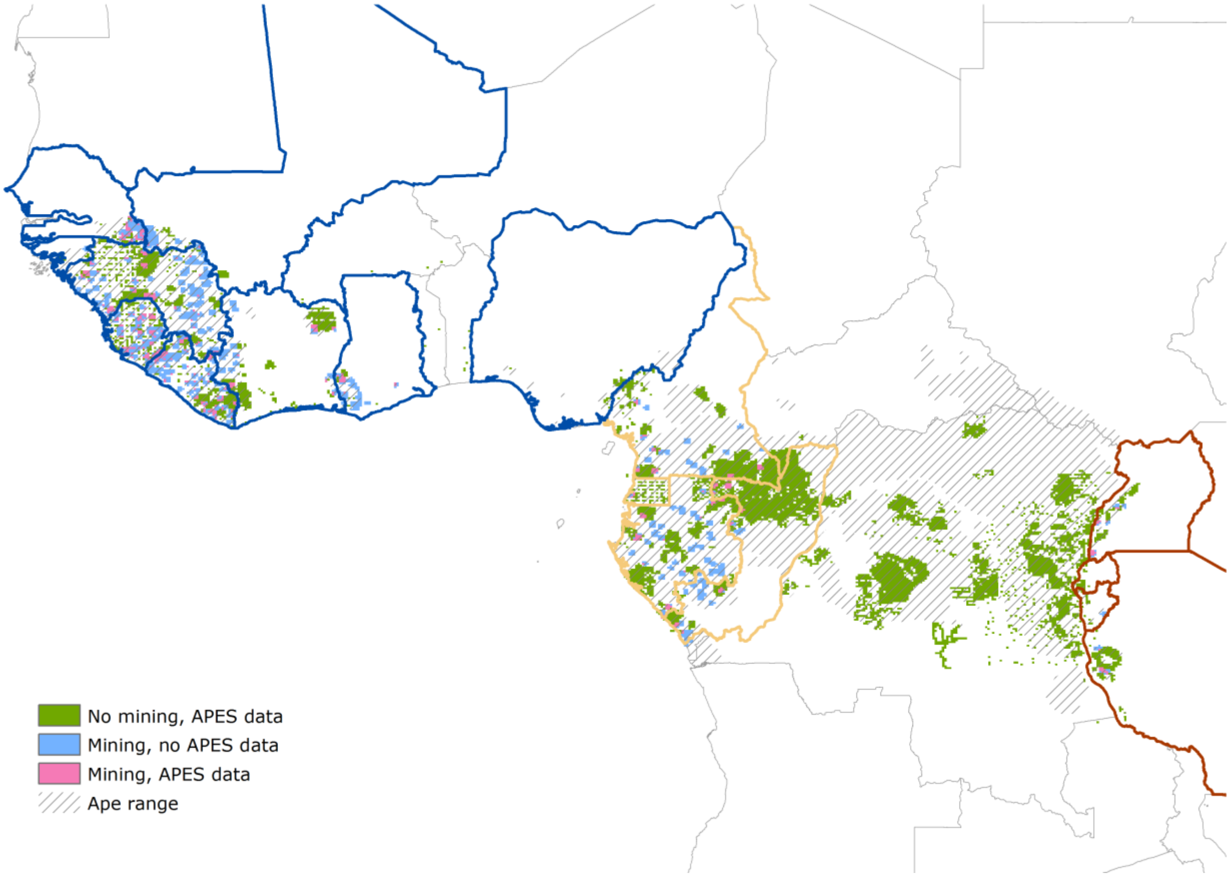


**Fig. S6.**

Distribution of areas surveyed for apes outside (green) and inside (pink) mining areas for which data have been archived in the IUCN SSC A.P.E.S. database. Areas that overlap with mining areas and for which no data currently exist in this database are marked in blue.

**Table S1.**

Summary statistics of the t-test iterations.

| Country | Buffer | Mining stage | Ape density inside mining area | Ape density outside mining area | Delta (density) | t | p |
|---|---|---|---|---|---|---|---|
| Burundi | 10 km | Pre-operational | 0.011638188 | 0.031590731 | -0.019952543 | -4.507.293.406 | 0.001517927 |
| Cameroon | 10 km | Operational | 0.180626455 | 0.176733369 | 0.003893085 | 0.450978239 | 0.473900012 |
| Cameroon | 10 km | Pre-operational | 0.166250598 | 0.176854398 | -0.0106038 | -4.144.231.793 | 0.001960298 |
| Congo | 10 km | Operational | 0.439514051 | 0.880613391 | -0.44109934 | -1.386.245.155 | 4.95E-29 |
| Congo | 10 km | Pre-operational | 0.426578742 | 0.881770027 | -0.455191284 | -3.544.981.685 | 5.71E-225 |
| Equatorial Guinea | 10 km | Pre-operational | 0.196844397 | 0.282826184 | -0.085981787 | -1.316.353.314 | 1.42E-26 |
| Gabon | 10 km | Operational | 0.420340257 | 0.459753729 | -0.039413471 | -2.692.699.409 | 0.051678722 |
| Gabon | 10 km | Pre-operational | 0.434007938 | 0.459644476 | -0.025636538 | -6.025.848.181 | 3.81E-06 |
| Ghana | 10 km | Operational | 0.001193925 | 0.000734062 | 0.000459863 | 2.293.726.831 | 0.088614268 |
| Ghana | 10 km | Pre-operational | 0.000805535 | 0.000727298 | 7.82E-05 | 0.715163054 | 0.422299934 |
| Guinea | 10 km | Operational | 0.082441709 | 0.117805202 | -0.035363493 | -1.018.947.128 | 4.90E-17 |
| Guinea | 10 km | Pre-operational | 0.082809034 | 0.117900886 | -0.035091852 | -2.315.112.192 | 9.11E-95 |
| Guinea-Bissau | 10 km | Pre-operational | 0.051680894 | 0.054745044 | -0.00306415 | -0.37573626 | 0.47403811 |
| Côte d'Ivoire | 10 km | Operational | 0.002953378 | 0.004491999 | -0.001538621 | -1.291.528.735 | 0.30270511 |
| Côte d'Ivoire | 10 km | Pre-operational | 0.00302172 | 0.004518158 | -0.001496437 | -4.476.388.942 | 0.001911229 |
| Liberia | 10 km | Operational | 0.068105182 | 0.068056538 | 4.86E-05 | 0.013861984 | 0.508277722 |
| Liberia | 10 km | Pre-operational | 0.072520975 | 0.068121629 | 0.004399346 | 5.074.631.434 | 0.000100875 |

| | | | | | | | |
|---|---|---|---|---|---|---|---|
| Mali | 10 km | Operational | 0.014976436 | 0.01796701 | -0.002990574 | -1.285.027.019 | 0.288851045 |
| Mali | 10 km | Pre-operational | 0.015547243 | 0.017863073 | -0.00231583 | -1.969.568.711 | 0.15093424 |
| Senegal | 10 km | Operational | 0.085338966 | 0.032824987 | 0.05251398 | 9.611.755.652 | 7.60E-10 |
| Senegal | 10 km | Pre-operational | 0.07220381 | 0.032830415 | 0.039373395 | 1.752.467.108 | 5.95E-33 |
| Sierra Leone | 10 km | Operational | 0.099492755 | 0.064999394 | 0.034493361 | 4.679.786.441 | 0.000278563 |
| Sierra Leone | 10 km | Pre-operational | 0.122393648 | 0.064960702 | 0.057432946 | 112.213.633 | 3.65E-24 |
| Tanzania | 10 km | Pre-operational | 0.018332528 | 0.01548143 | 0.002851098 | 1.700.079.348 | 0.206209531 |
| Uganda | 10 km | Operational | 0.050997944 | 0.03740852 | 0.013589424 | 1.554.099.701 | 0.233615488 |
| Uganda | 10 km | Pre-operational | 0.04448359 | 0.037428451 | 0.007055138 | 1.861.821.576 | 0.173787147 |
| Burundi | 50 km | Operational | 0.020020946 | 0.079737032 | -0.059716086 | -3.533.388.062 | 2.94E-190 |
| Burundi | 50 km | Pre-operational | 0.017535976 | 0.079776836 | -0.06224086 | -4.604.866.596 | 0 |
| Cameroon | 50 km | Operational | 0.196097456 | 0.167677364 | 0.028420093 | 1.244.741.187 | 1.35E-22 |
| Cameroon | 50 km | Pre-operational | 0.180568012 | 0.167736614 | 0.012831398 | 1.915.138.795 | 8.93E-64 |
| Congo | 50 km | Operational | 0.636668201 | 0.954520482 | -0.317852281 | -3.443.535.282 | 7.34E-211 |
| Congo | 50 km | Pre-operational | 0.598391753 | 0.954594492 | -0.356202739 | -6.290.485.326 | 0 |
| Equatorial Guinea | 50 km | Pre-operational | 0.252561301 | 0.291405234 | -0.038843933 | -1.925.305.047 | 6.06E-65 |
| Gabon | 50 km | Operational | 0.491196206 | 0.400262088 | 0.090934118 | 2.801.983.456 | 1.33E-139 |
| Gabon | 50 km | Pre-operational | 0.512947955 | 0.40020572 | 0.112742235 | 7.408.151.606 | 0 |
| Ghana | 50 km | Operational | 0.000885132 | 0.000931537 | -4.64E-05 | -0.131361736 | 0.497518613 |
| Ghana | 50 km | Pre-operational | 0.000645362 | 0.00092673 | -0.000281369 | -0.836891663 | 0.374650481 |
| Guinea | 50 km | Operational | 0.116337613 | 0.106260081 | 0.010077532 | 9.614.639.349 | 1.07E-14 |

| | | | | | | | |
|---|---|---|---|---|---|---|---|
| Guinea | 50 km | Pre-operational | 0.133996894 | 0.106280987 | 0.027715907 | 2.457.292.857 | 2.12E-113 |
| Guinea-Bissau | 50 km | Pre-operational | 0.068877147 | 0.045553776 | 0.023323371 | 107.969.938 | 8.74E-19 |
| Côte d’Ivoire | 50 km | Operational | 0.00419379 | 0.005308866 | -0.001115075 | -3.954.523.329 | 0.007234691 |
| Côte d’Ivoire | 50 km | Pre-operational | 0.004419231 | 0.005301041 | -0.000881811 | -4.243.695.947 | 0.002598921 |
| Liberia | 50 km | Operational | 0.068653479 | 0.04460189 | 0.02405159 | 1.129.831.694 | 2.53E-17 |
| Liberia | 50 km | Pre-operational | 0.088535102 | 0.044520997 | 0.044014106 | 2.004.524.744 | 5.34E-69 |
| Mali | 50 km | Operational | 0.01910823 | 0.010834707 | 0.008273523 | 4.674.740.154 | 0.001667356 |
| Mali | 50 km | Pre-operational | 0.025423923 | 0.01089057 | 0.014533353 | 7.674.559.163 | 4.30E-08 |
| Nigeria | 50 km | Operational | 0.000835946 | 0.005240337 | -0.004404391 | -1.180.587.183 | 3.25E-24 |
| Nigeria | 50 km | Pre-operational | 0.000672625 | 0.005238227 | -0.004565602 | -1.673.702.327 | 4.58E-50 |
| Rwanda | 50 km | Operational | 0.001843287 | 0.03957955 | -0.037736263 | -1.408.376.703 | 7.49E-30 |
| Rwanda | 50 km | Pre-operational | 0.000835882 | 0.039317781 | -0.038481899 | -4.961.543.593 | 0.000136427 |
| Senegal | 50 km | Operational | 0.077432203 | 0.012692115 | 0.064740088 | 3.020.343.618 | 3.37E-170 |
| Senegal | 50 km | Pre-operational | 0.113960046 | 0.012694311 | 0.101265734 | 1.253.280.545 | 4.28E-30 |
| Sierra Leone | 50 km | Operational | 0.067614929 | 0.167912479 | -0.10029755 | -1.383.089.143 | 1.18E-24 |
| Sierra Leone | 50 km | Pre-operational | 0.142967012 | 0.167882423 | -0.024915411 | -2.638.385.812 | 0.054320188 |
| Tanzania | 50 km | Pre-operational | 0.032257666 | 0.010193988 | 0.022063677 | 3.523.887.187 | 5.30E-226 |
| Uganda | 50 km | Operational | 0.051113241 | 0.02447856 | 0.026634681 | 1.567.325.982 | 1.18E-37 |
| Uganda | 50 km | Pre-operational | 0.045878346 | 0.024517811 | 0.021360536 | 1.795.297.512 | 2.57E-48 |

**Table S2.**

Total- and proportional overlap between ape density distribution and mining areas with 10 km- and 50 km buffers in the different range countries.

| Region | No. apes threatened by mining (10 km buffers) | Proportional overlap % (10 km buffers) | No. apes threatened by mining (50 km buffers) | Proportional overlap % (50 km buffers) |
|---|---|---|---|---|
| Tanzania | 58 | 3 | 988 | 57 |
| Rwanda | 0 | 0 | 29 | 14 |
| Burundi | 20 | 3 | 437 | 58 |
| Uganda | 214 | 5 | 2822 | 70 |
| Cameroon | 1292 | 2 | 23300 | 37 |
| Congo | 3683 | 1 | 55020 | 20 |
| Gabon | 5546 | 5 | 82005 | 69 |
| Equatorial Guinea | 62 | 1 | 1611 | 23 |
| Nigeria | 0 | 0 | 35 | 3 |
| Ghana | 25 | 45 | 56 | 100 |
| Côte d’Ivoire | 73 | 5 | 1125 | 80 |
| Mali | 356 | 19 | 1604 | 87 |
| Liberia | 1184 | 18 | 6505 | 99 |
| Guinea | 2036 | 7 | 23426 | 83 |
| Sierra Leone | 1341 | 26 | 4783 | 93 |
| Guinea-Bissau | 30 | 2 | 818 | 49 |
| Senegal | 623 | 27 | 1922 | 69 |
